# GenGraph: a python module for the simple generation and manipulation of genome graphs

**DOI:** 10.1101/465831

**Authors:** Jon Mitchell Ambler, Shandukani Mulaudzi, Nicola Mulder

## Abstract

**Background:** As sequencing technology improves, the concept of a single reference genome is becoming increasingly restricting. In the case of *Mycobacterium tuberculosis*, one must often choose between using a genome that is closely related to the isolate, or one that is annotated in detail. One promising solution to this problem is through the graph based representation of collections of genomes as a single genome graph. Though there are currently a handful of tools that can create genome graphs and have demonstrated the advantages of this new paradigm, there still exists a need for flexible tools that can be used by researchers to overcome challenges in genomics studies.

**Results:** We present the GenGraph toolkit, a tool that uses existing multiple sequence alignment tools to create genome graphs. It is written in Python, one of the most popular coding languages for the biological sciences, and creates the genome graphs as Python NetworkX graph objects. The conceptual model is highly intuitive, and as much as possible represents the biological relationship between the genomes. This design means that users will quickly be able to start creating genome graphs and using them in their own projects.

We outline the methods used in the generation of the graphs, and give some examples of how the created graphs may be used. GenGraph utilises existing file formats and methods in the generation of these graphs, allowing graphs to be visualised and imported with widely used applications, including Cytoscape, R, and Java Script.

**Conclusion:** GenGraph provides a set of tools for generating graph based representations of sets of sequences with a simple conceptual model in a widely used coding language. It is publicly available on Github (https://github.com/jambler24/GenGraph).

## Background

Modern genomics relies heavily on the use of a reference genome for common processes like variant calling, gene expression analysis, and even genome assembly. This reference sequence is often a consensus from a set of sequences that collectively represent anything from an individual isolate such as *Mycobacterium tuberculosis* H37Rv, to an entire species, in the case of the human genome assembly GRCh38, and the use of this single reference introduces a number of biases. The reference may be missing genes from some strains resulting in them being ignored in a differential expression analysis, or contain chromosomal rearrangements resulting in the effect of an upstream variant being misinterpreted. In terms of genome storage, the current standard is as linear sequence stored in a fasta file. Although they have served their purpose up until now, in the age of pangenomes and microbiome studies these representations have become limiting in terms of the file space they occupy, the functionality they provide, and their ability to represent population scale variation. These challenges have led to a move towards genome graphs. Where a pangenome is a collection of sequences that represent all variation between individuals in a defined clade, a genome graph is a graph representation of a pangenome where the sequences can be represented as a De Bruiijn graph, directed acyclic graph, bidirected graph, or a biedged graph. This representation of genomes offers a myriad of advantages over the use of a single reference genome.

Graphs are not a new concept in genomics, and are used for tasks including the assembly of genomes and the alignment of reads. Now, tools like vg (variant graph) [1], PanTools [2] and the Seven Bridges genome graph toolkit (https://www.sbgenomics.com/graph/) allow for the creation and utilisation of genome graphs in genomics work flows. These tools are developing rapidly, and include features that take advantage of the graph structure, allowing for read alignment and variant calling using the graph genome as a reference. While these tools are highly capable, there still exists a need for the development of more toolkits for genome graphs as advocated by Paten *et al*., in a recent review that discusses these tools and the improvements they have brought to variant calling [3].

GenGraph is a genome graph creation and manipulation toolkit created in Python that focuses on providing tools for working with bacterial genome graphs. It is able to create a genome graph using multiple whole genomes and existing multiple sequence alignment (MSA) tools, allowing any current or future algorithm to be employed. In this article we outline the structure of the genome graph created by GenGraph, the methods for its creation. Further examples of applications are available on the project Github page.

## 1 Implementation

GenGraph was implemented as both a Python tool and a module so that it may be used stand alone or as part of another tool. The graphs are NetworkX graph objects whose attributes may be accessed in the manner described in the NetworkX documentation.

### 1.1 Structure of the graph

GenGraph creates a directed sequence graph, where the individual genomes are encoded as walks within the graph along a labeled path (Figure 3D). Each node ideally represents a sub-sequence that is homologous between the component sequences. This implies the sequences have a shared evolutionary origin, and are not just identical sequences, and biological representation of the genomes is prioritised over a compressed data structure (Figure 1). This is an important design choice in that it allows for a more intuitive use of the graph, and a simpler conceptual model. As the graph contains no self loops, creating functions that require traversal is kept simpler. The coordinate system allows for existing annotations to be used, and for the sequence of a gene for a particular isolate to be retrieved from the graph given a traditional GTF file, a common task for which a function has been created. The coordinate system also allows for inversions to be represented in a single node (Figure 2). This results in a more intuitive representation of the relationship between the sequences as genes that fall within the inverted sequence are still found in the same node in both isolates, and the concept of a chromosomal breakpoint is represented by the edges either side of the node. Descriptions of the node and edge attributes can be found in tables 1 and 2.

**Figure 1.**
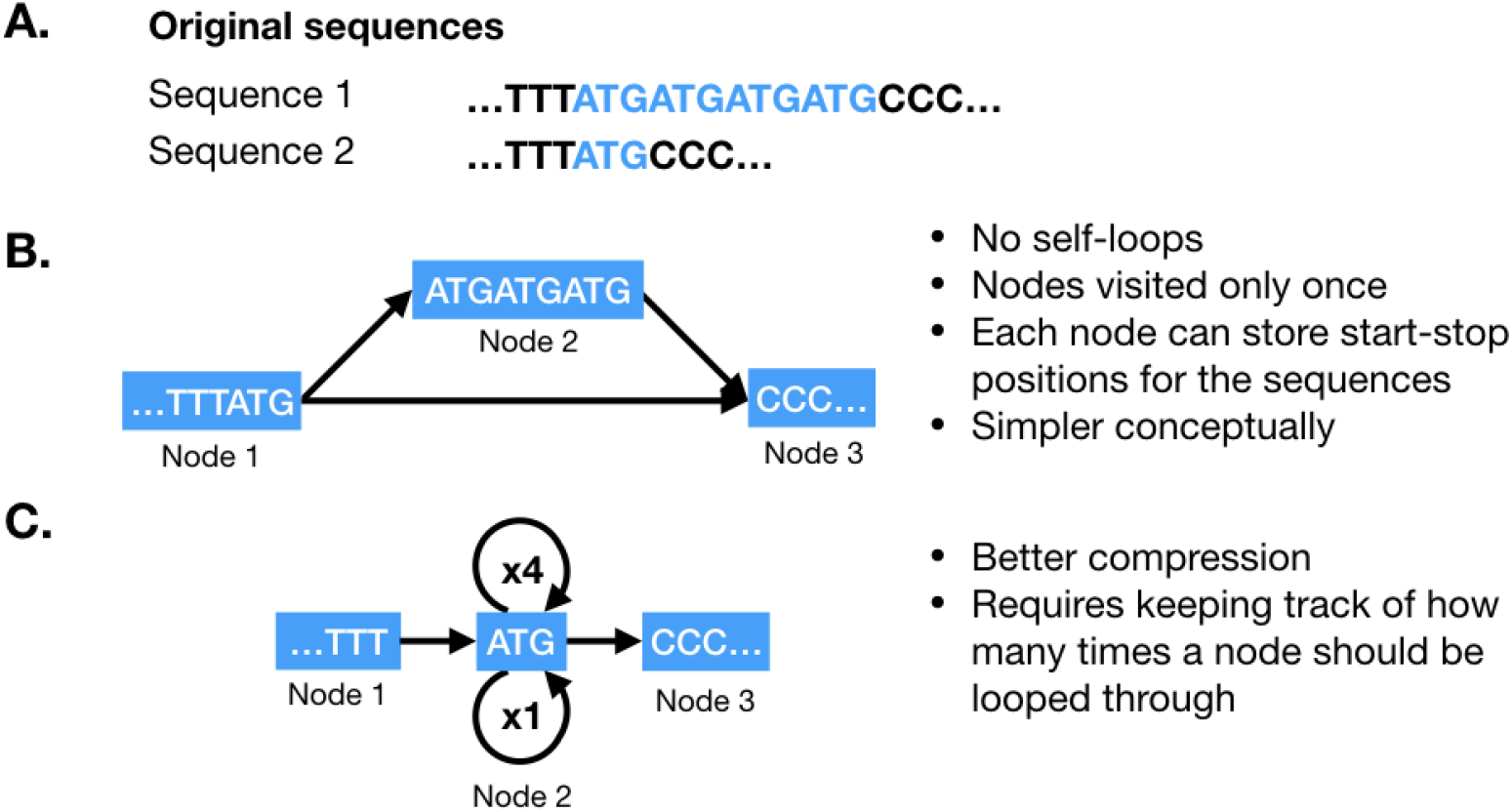
Representation of repeats in the genome graph. **A**, Two sequences where sequence 2 contains 3 additional “ATG” repeats high-lighted in blue. **B**, GenGraph represents only differences, with node 1 representing both sequences, node 2 representing the additional repeats found only in sequence 1, and node 3 the sequence that is once again shared. **C**, This is opposed to an approach where the “ATG” repeat is represented as a single node with a self loop. This approach may be neater and result in better compression, but raises many practical problems including not allowing the node to be labeled with the sequence start and stop positions.

**Figure 2.**
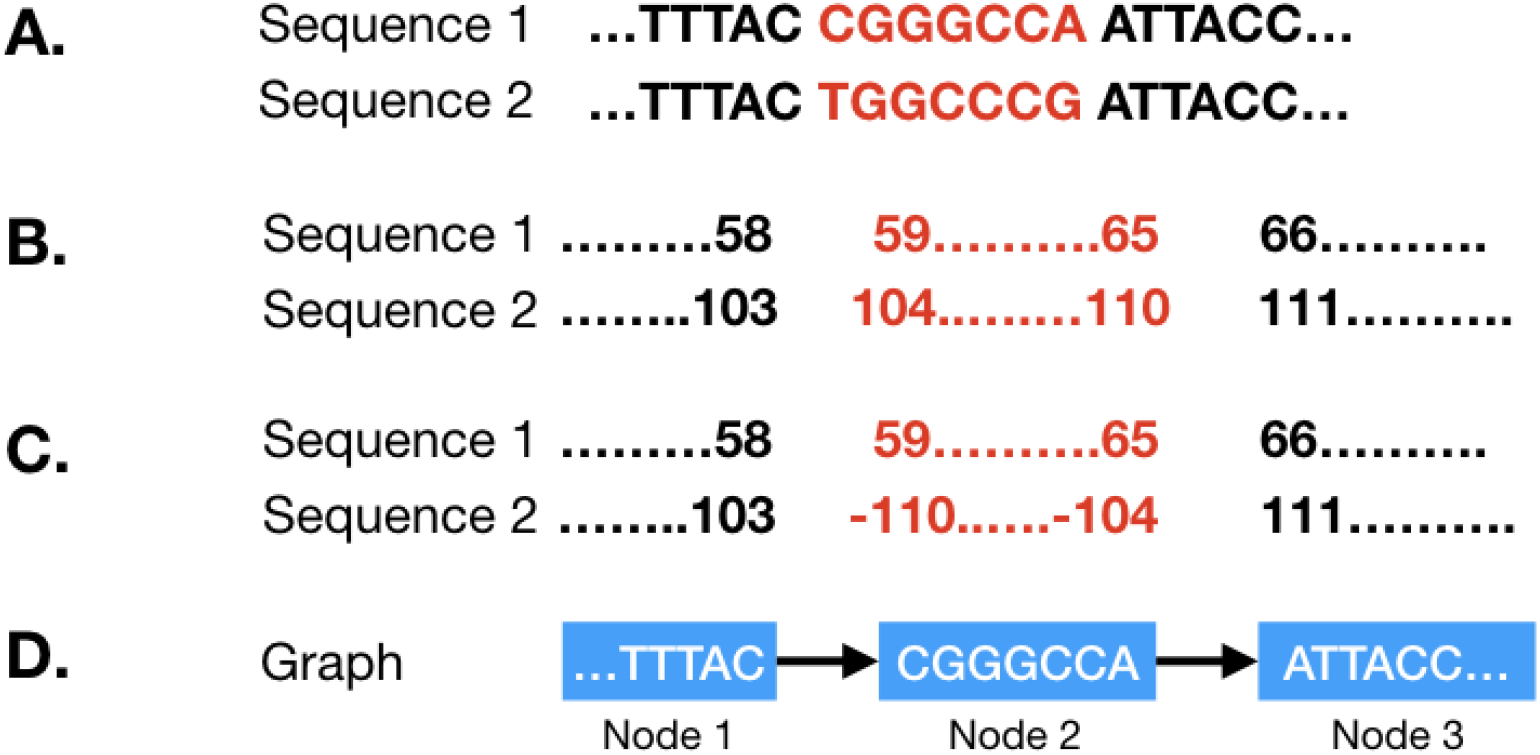
Representation of inversions in the genome graph. During the first step of genome graph creation, co-linear blocks are identified. In some cases, these may be homologous sequences that have been inverted. GenGraph represents these sequences in a single node (that may be broken down into more nodes in the second step) and represents the inverted state of the sequence by negative nucleotide position values in the node. **A**, Two sequences are shown where an inversion has taken place. This is normally a larger stretch of sequence perhaps a few kb in length. **B**, The positions of the sequences are different, as is generally the case with homologous sequences. The positions of the nucleotides flanking the breakpoints are shown. **C**, The inversion in the second sequence is represented by reversed negative nucleotide position values. **D**, This way, both sequences are represented in the same node, and to recreate sequence 2, the sequence in the node is simply reverse-complimented.

**Figure 3.**
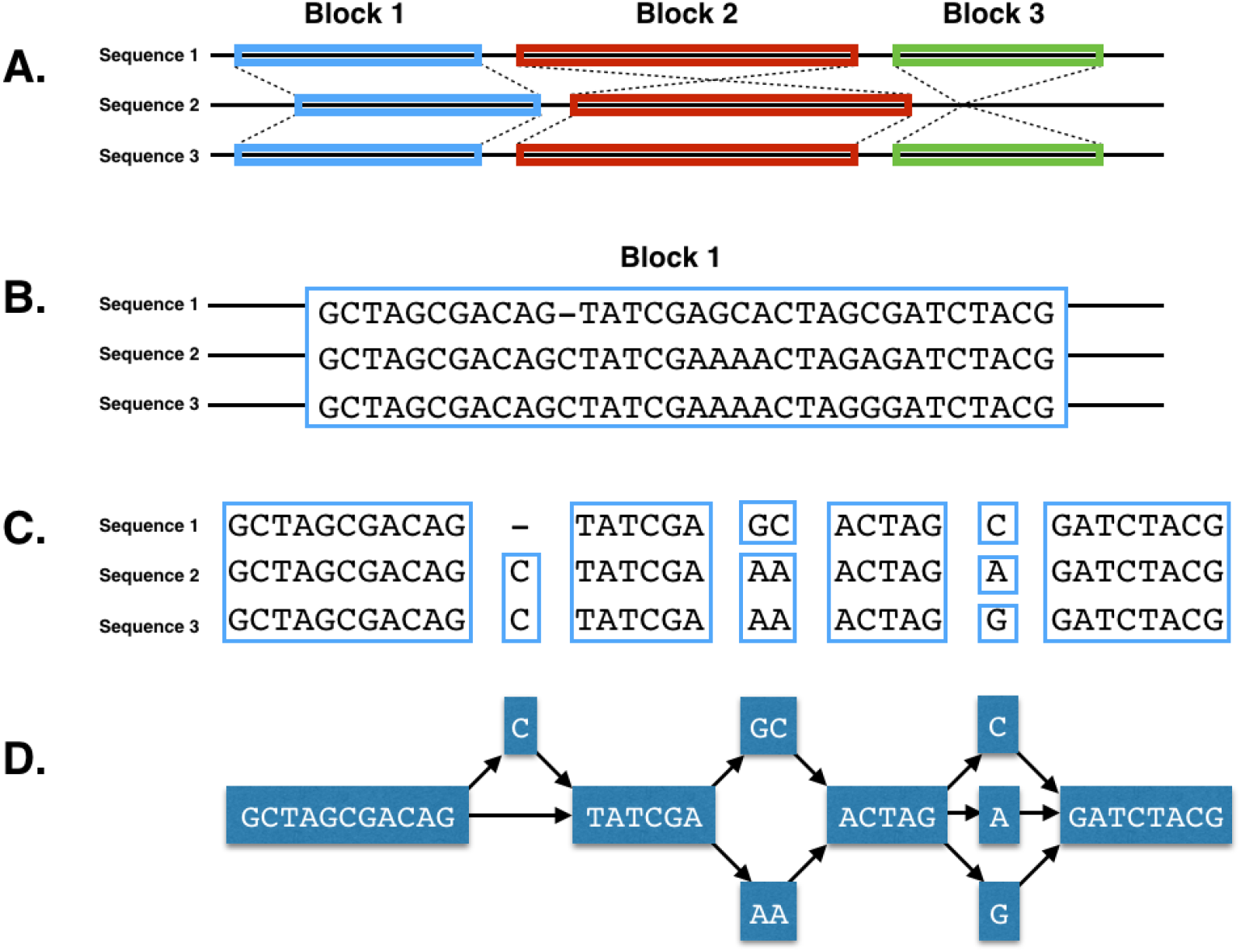
Overview of the GenGraph algorithm. **A**, Co-linear blocks of sequence are identified to determine the structural relationship of the sequences. **B-C**, Each block is then realigned using a MSA tool. **C-D**, Identical sequences are reduced into nodes and edges created.

**Table 1.** Information on the node attributes used in a gengraph genome graph.

| Node attribute | Type | Description |
| --- | --- | --- |
| name | String | A unique name identifying the node. When a node is split, the resulting nodes inherit the original node’s value appended with a new number. So if node Aln_66 is split into 4 nodes, they are named Aln_66_1, Aln_66_2, Aln_66_3, Aln_66_4. |
| sequence | String | The nucleotide sequence that is represented by this node. |
| ids | String | This is a comma separated list of the isolates that are represented by this node. |
| (isolate)_leftend,<br>(isolate)_rightend | Integer | For each isolate, the positions of the first and last nucleotides represented by the node is recorded. So for isolate H37Rv, H37Rv_leftend = 13,203. |

**Table 2.** Information on the edge attributes used in a gengraph genome graph.

| Edge attribute | Type | Description |
| --- | --- | --- |
| name | String | Edges are named by a combination of the two nodes that they link. Eg: Aln_48_50 (-) Aln_48_49 would be the name of the node that links nodes Aln_48_50 and Aln_48_49 |
| ids | String | This is a comma separated list of the isolates that are represented by this edge. |

### 1.2 Creation of the genome graph

The graph is created as a Python NetworkX graph object, the details of which may be found here (https://networkx.github.io/documentation/networkx-1.9/overview.html). GenGraph creates the genome graph in two steps (Figure 3). First, large structural differences between the genomes including large deletions and chromosomal inversions are identified by finding large blocs of co-linear sequence between the genomes using a tool like progressiveMauve [4]. These represent regions of structural conservation, and are temporarily stored in a single node within the initial graph, even if they are imperfect alignments. GenGraph then realigns the sequences in these initial nodes using the selected MSA tool, and finds the best local alignment for the sequences. The relative start and stop positions for the region of aligned sequence contained within the node is stored for each of the isolates that are included in that node as attributes of the NetworkX node object. This is then converted into a sub-graph by collapsing shared regions into single nodes, and creating edges so that a path exists for each of the original sequences through the sub-graph. This sub-graph then replaces the initial temporary node from the initial structural graph.

The process of identifying the co-linear blocks and subsequent realignment is done using functions that wrap existing alignment tools. Currently Muscle [5], Mafft [6] and Clustal Omega [7] are supported for the local MSA. The final NetworkX graph objects created by GenGraph may be exported as GraphML, XML, or as a serialised object,though various other formats may be added in future. GenGraph creates a report file containing information such as the number of nodes and edges in the graph, the average in and out degree of the nodes, the total sequence length of all the nodes in the graph and the density of the graph (Additional file 1). This information can be used to monitor how graphs change as more genomes are added as well as the relationship between the number of features and the graph size.

### 1.3 Novel graph functions

GenGraph is available as both a command line tool, and a Python module. Both allow for the creation of a genome graph, and the use of an existing genome graph for downstream analysis. These functions include simple processes like extracting a single genome in fasta format for a specific isolate, or extracting a sub-sequence. The functions can be used together to create new tools that use the genome graph as a reference. As examples, the GenGraph command line tool can be used to extract a pan-genome from the genome graph, or to identify homologous genes between isolates.

To demonstrate the use of GenGraph, we downloaded the complete genome assemblies of various *Mycobacterium tuberculosis* isolates from the NCBI database (https://www.ncbi.nlm.nih.gov/genome/) and used them to construct a genome graph using progressiveMauve for the structure graph, and Clustal Omega for the realignment of the blocks. This genome graph was then used to demonstrate the pan-genome extraction and gene homology functions.

## 2 Results and Discussion

### 2.1 Structure of the graph

GenGraph creates the genome graph based on whole genome sequence alignments that are conducted in two parts. First, the identification of large co-linear blocks, then the realignment of those blocks. This allowed for the creation of a genome graph containing multiple bacterial genomes, including the MTB isolate W-148 that contains large scale chromosomal rearrangements [9]. The created genome graph is thus able to capture all variants from large scale structural differences between isolates such as chromosomal rearrangements, down to smaller scale differences including SNPs and copy number variations.

The structure aims to represent a biologically accurate representation of the evolutionary relationships between the sequences of the different isolates. In doing so, it also maintains a simple conceptual model, which makes interpreting the graph simple, and in turn helps developers to create new tools easier.

### 2.2 Global and local alignment: Performance

A primary feature of GenGraph is the use of existing MSA tools for the creation of the graph by wrapping the tools in functions. Because GenGraph uses these alignment tools to create the graph structure, users may use parameters or aligners that are best suited for the organism. With MSA being the current speed bottleneck in the creation of genome graphs, the scalability of GenGraph is dependent on the ability of these alignment tools. This allows the toolkit to evolve and improve with time, as well as utilise alignment tools that are best suited to the dataset at hand, and adapt to new innovations including GPU acceleration or field-programmable gate array chips.

Graph generation runtime increases in a linear fashion, influenced by the number of sequences being aligned, their length, and their similarity. By breaking down the genomes into partially pre-aligned blocks, GenGraph is able to align multiple long genomes in segments and with the current version of mafft able to align up to 30,000 sequences in a block.

GenGraph was able to create a genome graph containing 5 MTB genomes on a 2012 i7 Macbook Pro with 8 GB ram using Mafft in 53 minutes and 10 genomes in 2 hours and 44 minutes (Additional file 2). For smaller genomes, 300 HIV-1 genomes were aligned and converted to a genome graph in 35 minutes. Currently GenGraph does not take advantage of multiprocessing, an enhancement that will be made in an upcoming update. From testing we see the scalability of GenGraph is dependent on the power of the latest alignment tools, the number of sequences being aligned, their length, and their similarity, though in general we observe a linear increase in genome graph generation time as the number of sequences increase (Table 3). Graph generation represents the most computationally intense and time consuming process, while downstream analysis benefits from the use of the data in an aligned form.

**Table 3.** An increase in file size was observed per genome added to the graph that demonstrated the compression of data that occurs by collapsing regions of shared aligned sequences into single representative nodes. The compression is related to the similarity of the sequences, as sequences that only differ by few bases will only require a few additional nodes.

| Number of genomes | 1 | 2 | 3 | 4 | 5 | 6 | 10 |
| --- | --- | --- | --- | --- | --- | --- | --- |
| File size | 4,5Mb | 5,9Mb | 7,6Mb | 8,5Mb | 11Mb | 13Mb | 38Mb |
| Number of nodes | 0 | 3,690 | 8,106 | 9,320 | 13,264 | 15,355 | 43,290 |
| Number of edges | 0 | 4,886 | 10,868 | 12,485 | 17,823 | 22,296 | 73,652 |

### 2.3 Toolkit

Python, along with R, are widely used in the sciences. Python is particularly easy to learn, powerful, and has numerous tools like Biopython and Numpy that come with most of the functions needed to rapidly develop new tools. NetworkX is one of those tools, providing a number of functions for working with networks and graphs right out of the box.

As GenGraph was developed with the challenges of working with bacterial genomes in mind, a number of functions that utilise the genome graph structure have been created to overcome those challenges. Many of these tools take advantage of the MSA derived graph structure, speeding up processes that usually would require the use of BLAST such as finding gene orthologues between isolates. An example is a tool for the creation of a homology matrix, which is useful for interpreting results from isolates with poor or missing annotations. In *Mycobacterium tuberculosis* isolates, due to the variance in genome size and composition between strains, as well as the limited availability of well annotated reference genomes, researchers often have to choose between aligning reads to the closest related genome or to the available reference genome for the organism. The homology matrix created by GenGraph allows mapping of gene IDs between isolates to identify orthologues in a gene-order aware manner. As orthologues are identified by their relative position within in the graph, and not by their sequence similarities, the correct orthologues of genes with multiple paralogues such as the PE/PPE genes in *M. tuberculosis* are identified.

Additional tools for cladogram construction, ancestral genome reconstruction, and pan-transcriptome extraction have been created that take advantage of the genome graph structure in a similar manner and are outlined on the GetGraph GitHub page under the Wiki, as well as in supplementary file 3 where usage examples are also given. Using existing graph formats allows exported sequence graphs to be visualised in commonly available programs such as Cytoscape [8] allowing for the visualisation of genomic features and regions of interest.

## 3 Conclusions

The GenGraph toolkit provides a simple method to create genome graphs using MSA tools. It is able to scale from small viral genomes to bacterial genomes on desktop computers, with further testing for large genomes already underway. GenGraph uses external alignment tools for the creation of the alignments used in generating the genome graph, making it able to utilise any existing or future alignment tools to boost its performance. Because only existing graph file formats are used, the graphs can be used and visualised by common tools, including Cytoscape and R. To facilitate adoption, GenGraph was created with bacterial genomes in mind, and includes a number of useful tools in order to facilitate adoption into work flows and overcome some of the challenges of working with poorly annotated strains by identifying orthologues between species with superior annotations and presenting them in the unified structure of a genome graph. These genome graphs can serve as a replacement for using a single fasta file representing a single isolate as a reference, increasing the number of reads that can be aligned, as well as the accuracy, as they are being aligned to the closest available isolate. Combined with an intuitive conceptual model and built for Python, one of the most widely used programming languages in the biological sciences, GenGraph provides a starting point for the development of a new generation of tools utilising genome graphs.

## 4 Abreviations

GFF: General feature format.
GTF: Gene transfer format.
MSA: Multiple sequence alignment.
MTB: *Mycobacterium tuberculosis*.
SNP: Single nucleotide polymorphism.
SRA: Sequence Read Archive.
VCF: Variant call format.

## Ethics approval and consent to participate

Not applicable.

## Consent for publication

Not applicable.

## Availability of data and material

–Project name: GenGraph
–Project home page: https://github.com/jambler24/GenGraph
–Operating system(s): Linux, Mac, Windows
–Programming language: Python
–Other requirements: NetworkX, Mauve, and Mafft.
–License: GNU LGPL

Datasets used during the current study are available from the NCBI https://www.ncbi.nlm.nih.gov.

## Competing interests

The authors declare that they have no competing interests.

## Funding

This work was funded by the National Research Foundation of South Africa, grant number 86934.

## Author’s contributions

JMA, PhD student, writing, developing the code for GenGraph; NM, corresponding author, technical supervision and assistance, proof reading, Bioinformatics component; SM, development of GenGraph functions.

## Acknowledgements

We thank the National Research Foundation of South Africa for their financial support of this research, as well as the University of Cape Town for use of their facilities.

## Additional Files

**Additional file 1 — Example report file for the Genome graph created by GenGraph**

The report file is in .txt format and contains details of the generated genome graph.

**Additional file 2 — Genome graph containing five MTB genomes**

The graph was created from five MTB isolates (W148, F11, CDC1551, H37Rv, H37Ra, CCDC5180) and saved in GraphML format. This file is viewable in Cytoscape and may be imported using Python’s NetworkX package.

## References

1. VG Team: Variant Graph. https://github.com/vgteam/vg/

2. Sheikhizadeh, S., Schranz, M.E., Akdel, M., de Ridder, D., Smit, S.: PanTools: representation, storage and exploration of pan-genomic data. Bioinformatics 32(17), 487–493 (2016). doi:10.1093/bioinformatics/btw455

3. Paten, B., Novak, A.M., Eizenga, J.M., Garrison, E.: Genome graphs and the evolution of genome inference. Genome Res. 27(5), 665–676 (2017). doi:10.1101/gr.214155.116

4. Darling, A.E., Mau, B., Perna, N.T.: Progressivemauve: Multiple genome alignment with gene gain, loss and rearrangement. PLoS One 5(6) (2010). doi:10.1371/journal.pone.0011147

5. Edgar, R.C.: MUSCLE: Multiple sequence alignment with high accuracy and high throughput. Nucleic Acids Res. 32(5), 1792–1797 (2004). doi:10.1093/nar/gkh340

6. Katoh, K., Kuma, K.I., Toh, H., Miyata, T.: MAFFT version 5: Improvement in accuracy of multiple sequence alignment. Nucleic Acids Res. 33(2), 511–518 (2005). doi:10.1093/nar/gki198

7. Sievers, F., Wilm, A., Dineen, D., Gibson, T.J., Karplus, K., Li, W., Lopez, R., McWilliam, H., Remmert, M., Söding, J., Thompson, J.D., Higgins, D.G.: Fast, scalable generation of high-quality protein multiple sequence alignments using Clustal Omega. Mol. Syst. Biol. 7(1), 539 (2011). doi:10.1038/msb.2011.75

8. Shannon, P., Markiel, A., Ozier, O., Baliga, N.S., Wang, J.T., Ramage, D., Amin, N., Schwikowski, B., Ideker, T.: Cytoscape: A Software Environment for Integrated Models of Biomolecular Interaction Networks. Genome Res. 13(11), 2498–2504 (2003). doi:10.1101/gr.1239303

9. Shitikov, E.A., Bespyatykh, J.A., Ischenko, D.S., Alexeev, D.G., Karpova, I.Y., Kostryukova, E.S., Isaeva, Y.D., Nosova, E.Y., Mokrousov, I.V., Vyazovaya, A.a., Narvskaya, O.V., Vishnevsky, B.I., Otten, T.F., Zhuravlev, V.I., Zhuravlev, V.Y., Yablonsky, P.K., Ilina, E.N., Govorun, V.M.: Unusual large-scale chromosomal rearrangements in Mycobacterium tuberculosis Beijing B0/W148 cluster isolates. PLoS One 9(1), 84971 (2014). doi:10.1371/journal.pone.0084971

